# Motor reattachment kinetics play a dominant role in multimotor-driven cargo transport

**DOI:** 10.1101/180778

**Authors:** Qingzhou Feng, Keith J. Mickolajczyk, Geng-Yuan Chen, William O. Hancock

**Affiliations:** Department of Biomedical Engineering, Penn State University, University Park, Pennsylvania, USA; Intercollege Graduate Degree Program in Bioengineering, Penn State University, University Park, Pennsylvania, USA; Molecular Cellular and Integrative Biological Sciences Program in Huck Institute of Life Sciences, Penn State University, University Park, Pennsylvania, USA

**Keywords:** kinesin, microtubule, axonal transport, high frequency tracking, DNA origami

## Abstract

Kinesin-based cargo transport in cells frequently involves the coordinated activity of multiple motors, including kinesins from different families that move at different speeds. However, compared to the progress at the single-molecule level, mechanisms by which multiple kinesins coordinate their activity during cargo transport are poorly understood. To understand these multi-motor coordination mechanisms, defined pairs of kinesin-1 and kinesin-2 motors were assembled on DNA scaffolds and their motility examined *in vitro*. Although less processive than kinesin-1 at the single-molecule level, addition of kinesin-2 motors more effectively amplified cargo run lengths. By applying the law of total expectation to cargo binding durations in ADP, the kinesin-2 microtubule reattachment rate was shown to be 4-fold faster than that of kinesin-1. This difference in microtubule binding rates was also observed in solution by stopped-flow. High-resolution tracking of gold-nanoparticle-labeled cargo with 1 ms and 2 nm precision revealed that kinesin-2 motors detach and rebind to the microtubule much more frequently than do kinesin-1. Finally, cargo transported by kinesin-2 motors more effectively navigated roadblocks on the microtubule track. These results highlight the importance of motor reattachment kinetics during multi-motor transport and suggest a coordinated transport model in which kinesin-1 motors step effectively against loads while kinesin-2 motors rapidly unbind and rebind to the microtubule. This dynamic tethering by kinesin-2 maintains the cargo near the microtubule and enables effective navigation along crowded microtubules.

## INTRODUCTION

Kinesin motor proteins transport a diverse array of cargos to specific destinations in cells. One feature that helps to specify particular cargo to specific cellular locations is the spatial diversity of tubulin post-translational modifications and microtubule associated proteins (MAPs), with different kinesins walking preferentially on particular subsets of microtubules (1). Importantly, transport in axons and dendrites is generally bidirectional; hence cargo have both plus-ended kinesin motors and minus-ended dynein motors attached (2). Adding to this complexity, specific cargo can have two classes of kinesins simultaneously bound; for instance, synaptotagmin-rich axonal vesicles were shown to be transported simultaneously by kinesin-1 and kinesin-2 motors (3). Thus, to understand how specific cargo are targeted to specific locations in axons and dendrites, it is important to understand how motors coordinate their activities during multi-motor transport.

Because kinesin-1 and kinesin-2 motors move with two-fold different speeds in the absence of load (4), they do not appear to be an optimal pair for co-transport of intracellular cargos. They differ in other ways as well – compared to kinesin-1, heterotrimeric kinesin-2 motors are less processive and they detach much more readily under load (4-9). In contrast, kinesin-2 stepping is less affected than kinesin-1 by roadblocks on microtubules such as MAPs (10). A comprehensive understanding of bidirectional transport in neurons, and the transport defects that underlie neurodegenerative disease requires understanding both how uniform populations motors coordinate their transport activities and how diverse motors attached to a single cargo compete and coordinate to target cargo to their proper intracellular locations.

Although single kinesin-1 motors are robust transporters, previous experimental and theoretical work has suggested that they do not coordinate their activities well (11, 12). This property contrasts with dyneins – the finding that cargo stall forces are integer multiples of the single-dynein stall force has been used to argue that dyneins efficiently couple their activities during multi-motor transport (13, 14). The ability of different motors to coordinate their activities depends on their inherent unloaded velocity and directionality, as well as their ability to generate force and remain bound under load; properties that have been investigated extensively in single-motor experiments (2, 4, 6). In contrast, the rate that detached motors reattach to the microtubule during multi-motor transport is an equally important but understudied parameter. The importance of reattachment kinetics can be appreciated by taking the limits: if motor reattachment is instantaneous then all motors will be contributing to the transport at all times; whereas if motor reattachment is very slow then cargo movements are carried out by only one motor at any given time. Because experiments to date generally follow cargo position, rather than the dynamics of individual motors in a population, this reattachment rate is very difficult to determine experimentally, and in any case, it is expected to vary with the geometry of the cargo and motor-cargo linkages. Experiments with kinesin-driven membrane tethers estimated a reattachment rate of 4.7 s^-1^ in that particular geometry (15), and in modeling work, a reattachment rate of 5 s^-1^ has been used extensively for all kinesin and dynein isoforms (7, 16, 17). However, how this parameter varies for different motors and in different geometries is not clear.

The goal of the present work is to compare the degrees to which kinesin-1 and kinesin-2 coordinate their activities during multi-motor transport. In particular, we focus on the motor reattachment rate, and we find that kinesin-2 has a four-fold faster reattachment rate than kinesin-1. This finding suggests a multi-motor coordination scheme in which kinesin-1 provides sustained loads during long-distance transport and reattaches only slowly once it dissociates from the microtubule, while kinesin-2 frequently detaches and rapidly reattaches to the microtubule. This fast reattachment enables kinesin-2 to more efficiently explore the local microtubule landscape in cells and overcome roadblocks on microtubules such as MAPs and other cargos that may impede transport.

## MATERIALS AND METHODS

### Protein purification

Kinesin-1 assemblies consisted of *Drosophila* KHC truncated at 559 and fused to a C-terminus eGFP and His6 tag (4). Kinesin-2 consisted of the head and 17 amino acid neck-linker domain of *M. musculus* KIF3A fused to the coiled-coil of *Drosophila* KHC followed by eGFP and His6 tag, as previously described (4). Motors were bacterially expressed, purified by Ni column, and stored at -80 °C, following previously published protocols (4). For high-resolution tracking experiments, N-term biotinylated kinesin-1 and kinesin-2 motors were generated and attached to streptavidin-coated 30 nm gold nanoparticles (BBI Solutions) as previously described (18). Tubulin was purified from bovine brain as described (4). SNAP-tagged, His6-tagged GFP nano-body (GBP) (a gift from the Grischuck lab, University of Pennsylvania) was bacterially expressed and purified following protocols developed for motors (4).

### Generating oligo-functionalized GBP

Benzylguanine (BG) functionalized oligonucleotides were generated by reacting Benzylguanine-GLA N-hydroxysuccinimide (New England BioLabs) with C6-amine modified oligonucleotides (BG-oligo 1 and BG-oligo 2; **Fig. S1A**) in a 50 mM HEPES pH 8.5 buffer(19) for 30 mins, followed by purification through a Sephadex G-25 Superfine desalting column (GE Healthcare). BG oligos were then mixed with SNAP-tagged GBP for 1 h at 4 °C, followed by purification through the Ni column to remove un-reacted BG-oligos. GBP1 and GBP2 concentrations were quantified by mixing with varying known concentrations of complementary strands and running on SDS-PAGE gels to determine the concentration needed to completely shift the band to the higher molecular weight (**Fig. S1B**).

### Single molecule experiments

DNA scaffolds were labeled with Qdots (ThermoFisher) or gold nanoparticle (BBI Solutions). Motility solutions containing DNA scaffolds, oligo-functionalized GBP, motors, ATP or ADP were diluted in BRB80 (80 mM PIPES, 1 mM EGTA, 1 mM MgCl_2_, pH 6.8) to single molecule range (5 nM to 100 pM) with taxol, casein, BSA and antifade components described previously(4, 20). Taxol-stabilized microtubules were adsorbed onto cover slips of flow cells blocked with 2 mg/ml casein, motility solution introduced, and DNA scaffolds imaged by total internal reflection fluorescence microscopy (TIRFM) using a Nikon TE2000 microscope (60x, 1.45 NA PlanApo). Experiments were carried out at 21-23°C. Images were captured using a Cascade 512 CCD camera (Roper Scientific, Tucson, AZ) controlled by MetaVue software (Molecular Devices Corporation, Downingtown, PA). Run lengths and durations were analyzed by ImageJ (MTrackJ) using a pixel size of 71.0 nm. Kymographs were generated using Kymo-analyzer package (21). To ensure that run lengths were reliably captured, only run lengths greater than 200 nm were analyzed, and to estimate the average run length, this minimum distance was subtracted from all runs. High-resolution tracking methods and the associated image processing followed previously described protocols (18, 22).

### Data analysis

Mean and 95% confidence interval for run lengths and microtubule binding durations were estimated by Bootstrapping using MATLAB (Mathworks). Every data set was resampled with replacement 100 times, and generated data were fit to the exponential CDF 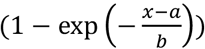 Reported mean and 95% confidence intervals were then calculated from the 100 resampled data sets (23).

Standard errors for kin1 and kin2 reattachment rates were calculated using the Error Propagation method (24). From Equation2,

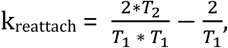

the standard error was calculated as

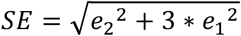

where e_1_ and e_2_ are the percent error of T_1_ and T_2_, respectively.

### Stopped-flow experiment

Stopped-flow experiments were carried out in BRB80 buffer in 23°C as previously described (25).

### Roadblock experiments

Microtubules with varying densities of roadblocks were made by polymerizing microtubules using varying ratios of biotinylated and unlabeled tubulin, incubating with saturating concentrations of neutravidin, and pelleting and resuspending to remove excess neutravidin. Total tubulin concentration was measured by A_280_ nm, and biotin concentration was measured using the HABA assay (Thermo Scientific).

## RESULTS

### Two-motor kinesin-2 assemblies have longer run lengths than two-motor kinesin-1 assemblies

To investigate defined teams of kinesin-1 and kinesin-2 motors, a SNAP-tagged anti-GFP nanobody (<u>G</u>FP <u>b</u>inding <u>p</u>rotein, GBP, 1 nM K_D_ for GFP (26) was used to link GFP-labeled motors to a quantum dot-functionalized DNA scaffold (**Fig. 1A**). Scaffolds containing either one or two motors were created by incubating scaffolds and free motors with either one or both GBP adapters (shown by gel in **Fig. 1B**). The kinesin-1 and -2 motors, which were fully characterized in previous work (4, 6, 27), share an identical coiled-coil domain and only differ by their motor domains, thus avoiding uncertainties regarding the effect of tether length or tail structure on motor behavior. One- and two-motor run lengths measured by total internal reflection fluorescence microscopy (TIRFM) were 0.77±0.16 μm and 1.62±0.23 μm, respectively for kinesin-1 (**Fig. 1C; Table S1**). The corresponding kinesin-2 run lengths were 0.65±0.13 μm and 2.38±0.26 μm (**Fig. 1D; Table S1**). Thus, adding a second motor increased the kinesin-1 run length by 2.1-fold and the kinesin-2 run length by 3.7-fold. Because a scaffold carried by two motors will continue to move as long as at least one motor is bound to the microtubule, the observed run lengths arise from two factors: the load-dependent detachment kinetics of each motor, and the rebinding rates of cargo-bound motors that has dissociated from the microtubule. A previous optical trapping study showed that the detachment rate of kinesin-2 is considerably more force dependent than that of kinesin-1 (6). Thus, the enhanced two-motor kinesin-2 run length suggests that the kinesin-2 reattachment kinetics are considerably faster than those of kinesin-1.

**Figure 1:**
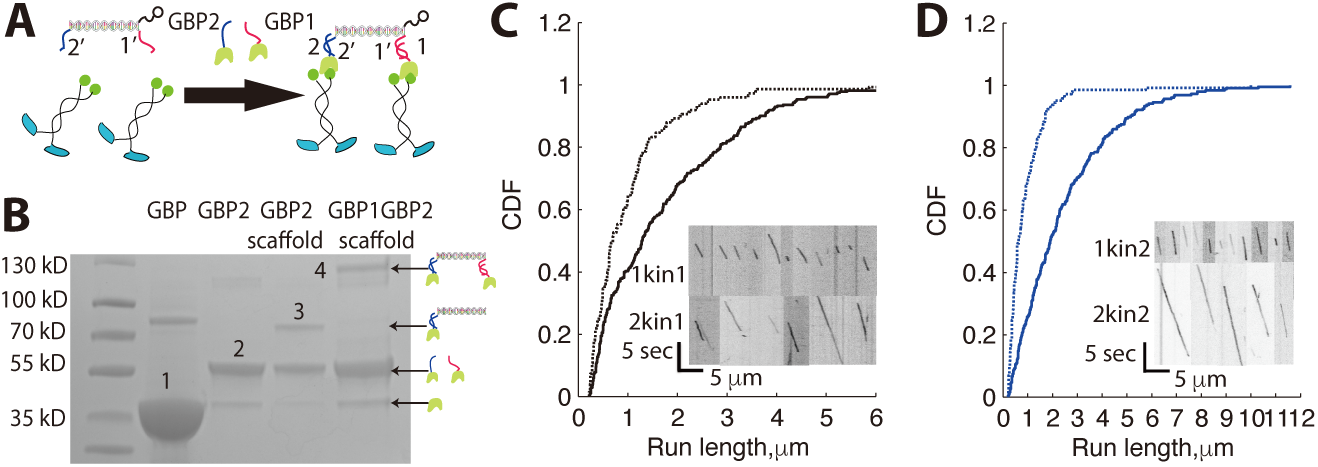
Assembly of defined multi-motor assemblies using DNA scaffold. (A) (Top) Schematic of DNA-motor assemblies. GFP binding proteins GBP1 and GBP2 were generated by covalently linking oligos 1 and 2 to the GBP through a C-terminal SNAP tag. GFP-labeled motors were then linked to the DNA scaffolds via overhanging single-stranded 1’ and 2’ appendages on the scaffold. Scaffolds were tracked by linking nano-particles to a third overhanging ssDNA on the scaffold. (B) SDS-PAGE gel of DNA-protein assemblies. Electrophoresis was performed on a 4% to 20% polyacrylamide gel. Labeled bands are: (1) unreacted GBP; (2) oligo-labeled GBP; (3) scaffold with one GBP bound; (4) scaffold with two GBP bound. ∼80 kD band in GBP lane is minor impurity from Ni-column purification. (C) Run lengths for assemblies containing one (dashed line) or two (solid line) kinesin-1 motors in 3 mM ATP, presented as cumulative distributions. Biotin-labeled scaffolds were mixed with GBP1 and excess motors to generate one-motor assemblies, and with both GBP1 and GBP2 to generate two-motor assemblies (**Fig. S1 C,D**). Inset: Kymographs of one-motor (upper) and two-motor (lower) runs for kinesin-1. Mean run lengths were 0.77±0.16 μm and 1.62±0.23 μm (mean±95% confidence interval, N=150 and N=283) for scaffolds containing one or two kinesin-1 motors, respectively. (D) Distribution and kymographs (inset) of kinesin-2 run lengths for one- (dashed line) and two- (solid line) kinesin-2 assemblies in 3 mM ATP. Mean run lengths were 0.65±0.13 μm (N=145) and 2.38±0.26 μm (N=257) for scaffolds containing one or two kinesin-2 motors, respectively.

### Kinesin-2 has a faster reattachment rate than kinesin-1

A parameter that, to our knowledge, has never been measured in a multi-motor complex is the rate that a dissociated motor bound to a cargo reattaches to the microtubule. To test our hypothesis that kinesin-2 has a faster reattachment rate than kinesin-1, we measured the binding duration of one- and two-motor assemblies in saturating ADP. In ADP, motors only bind and do not generate force, enabling us to make the important assumption that the detachment rate of each individual motor in a two-motor construct is the same as the detachment rate of one motor in ADP. Furthermore, because ADP release is the rate limiting step in solution (28), motors are initially in the ADP state upon microtubule binding independent of the nucleotide in solution, thus k_reattach_ measured in ADP should be the same as that in ATP. Using this approach, measured binding durations were interpreted using the model shown in **Fig. 2A**. Using TIRFM similar to the run length experiments, the mean one- and two-motor microtubule binding durations in ADP were T_1_= 0.72±0.15 s and T_2_=1.86±0.31 s for kinesin-1 and T_1_=0.50±0.11 s and T_2_=2.51±0.39 s for kinesin-2 (**Fig. 2B and C; Table S1**).

**Figure 2:**
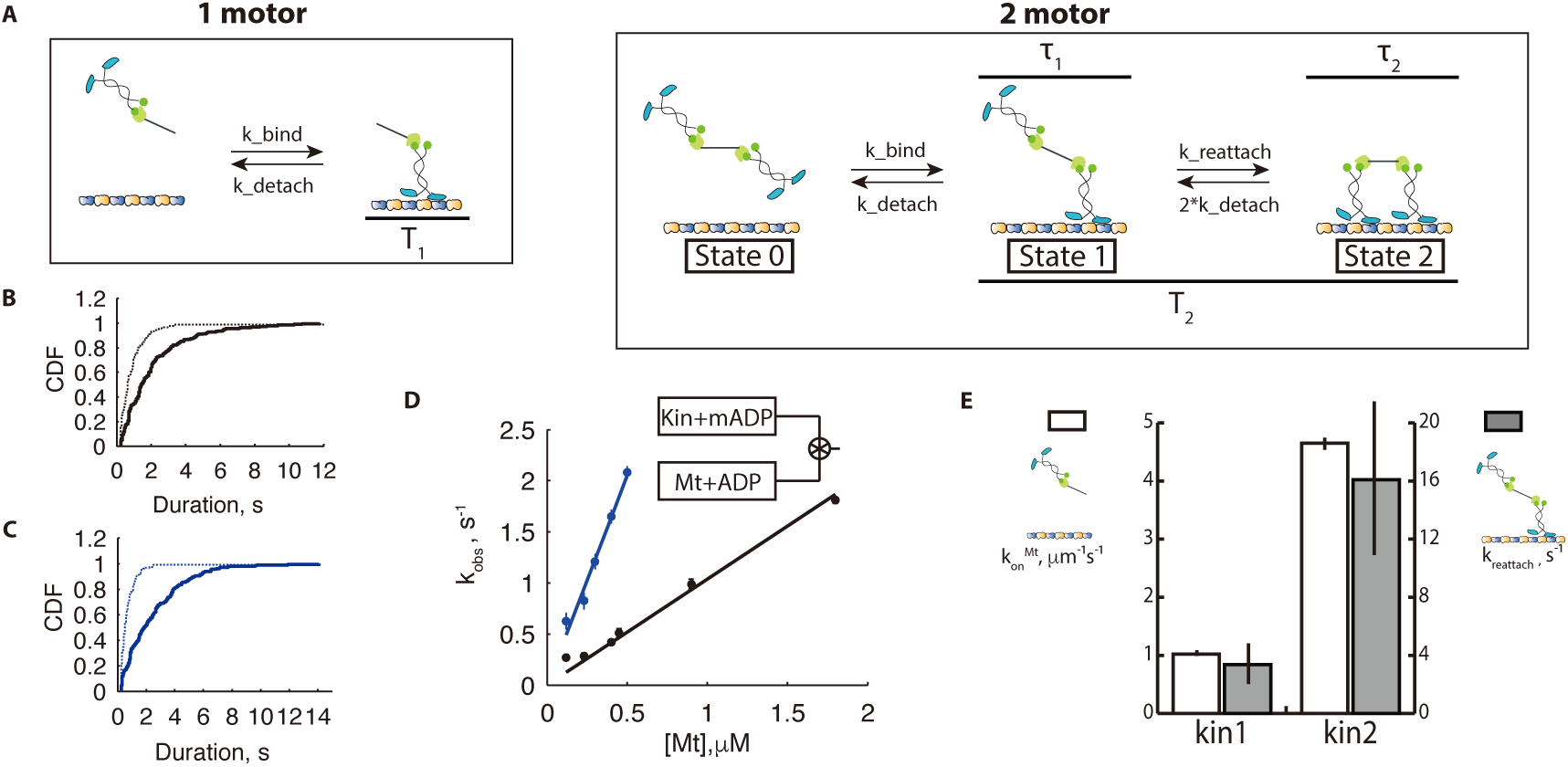
Calculating kinesin-1 and kinesin-2 reattachment rates from microtubule binding durations of one and two-motor assemblies in ADP. (A) Models used to analyze microtubule-binding durations. For one-motor assemblies (left), the microtubule binding duration in ADP (T_1_) is governed solely by the unbinding rate constant, k_detach_. For two-motor assemblies (right), a second parameter, the reattachment rate constant (k_reattach_) is introduced and an expression is derived for the expected two-motor binding duration in ADP (T_2_). See text for derivation. (B) Distribution of one- (dashed line) and two- (solid line) motor binding durations for kinesin-1 in 3 mM ADP. Mean binding durations were 0.72±0.15 s (N=90) and 1.86±0.31 s (N=223) for scaffolds containing one and two kinesin-1, respectively. See **Fig. S2A** for example kymographs. (C) Distribution of one- (dashed line) and two– (solid line) motor binding durations for kinesin-2 in in 3 mM ADP. Mean binding durations were 0.50±0.11 s (N=128) and 2.51±0.39 s (N=213) for scaffolds containing one and two-motor kinesin-2, respectively. See **Fig. S2B** for example kymographs. (D) Bimolecular on-rates for microtubule binding measured by stopped-flow. Observed motor binding rates were measured by fitting exponentials to the mantADP signal decay at varying microtubule concentrations (**Fig. S2 E**). Fitting a line to the measured rates at limiting [Mt] gives the bimolecular on-rate for microtubule binding k_on_Mt. Calculated k_on_Mt were 1.1±0.05 μM-1s^-1^ (regression ± RMSE) for kinesin-1 (black symbols) and 4.6±0.10 μM-1s^-1^ for kinesin-2 (blue symbols) **(Fig. S2 E, F)**. (E) Comparing bimolecular on-rates in solution to microtubule reattachment rates on scaffolds. Second-order k_on_MT (left axis, open bars from Fig. 2D) is 4.2-fold higher for kinesin-2 than kinesin-1. Similarly, the calculated first-order k_reattach_ (right axis, grey bars) is 3.6-fold faster for kinesin-2 than kinesin-1.

For one motor, the measured mean binding duration, T_1_ is simply the inverse of the first-order detachment rate, k_detach_. Thus, in saturating ADP, k_detach_ = 1.38±0.29 s^-1^ for kinesin-1 and k_detach_ = 2.00±0.44 s^-1^ for kinesin-2. For a two-motor complex, the observed binding duration, T_2_ includes states having either one or both motors attached. The importance of the k_reattach_ parameter is clear from inspection – a fast reattachment rate minimizes the probability that the complex is attached to the microtubule by only one motor, and hence minimizes the rate of detachment of the complex from the microtubule.

Based on the law of total expectation (29), we can calculate the reattachment rate for each motor from the measured T_1_ and T_2_, as follows. Starting from the initial state with one motor bound to the microtubule, there are two possibilities – either that motor will detach, terminating the event, or the second motor will attach to the microtubule. If the second motor attaches, then the complex will reside in a two-motor-bound state (state 2 in **Fig. 2A**) until either motor detaches, returning to the initial one-motor-bound state (state 1 in **Fig. 2A**). Because the system is memoryless, the duration starting from this revisited one-motor-bound state (state 1) is T_2_, just as before. Hence, if τ_1_ is the duration spent in state 1, τ_2_ is the duration spent in state 2, and P_12_ is the probability of the second motor binding (rather than the first motor dissociating), then the total binding duration can be calculated as

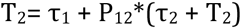

In this equation, the duration spent in state 1 is controlled by two transitions:

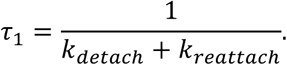

The duration spent in state 2 is:

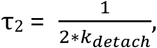

where the factor 2 is due to the fact that either motor can unbind, each with a rate k_detach_. Finally, the probability of the second motor binding (rather than the first motor detaching) is:

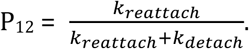

Solving for T_2_ (**Fig. 2A**), we get

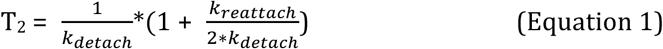

Solving for k_reattach_ in terms of the measured T_1_ and T_2_:

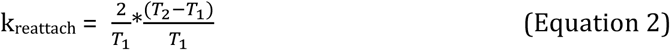

Plugging in the measured binding durations from **Fig. 2B and C**, k_reattach_ =4.41±1.75 s^-1^ for kinesin-1 and k_reattach_ =16.1±6.6 s^-1^ for kinesin-2, indicating that the reattachment rate of kinesin-2 is 3.6-fold faster than kinesin-1. To validate our result, we varied k_detach_ by lowering the level of ADP to 10 μM, which causes the motor to reside in the tight-binding apo state for a larger fraction of time, and repeated the analysis (**Fig. S2 C, D; Table S1**). This independentexperiment, which generated different T_1_ and T_2_ durations, resulted in similar k_reattach_ values of 4.6±3.2 s^-1^ for kinesin-1 and 18.7±8.0 s^-1^ for kinesin-2. This agreement supports the validity of the measurement and additionally confirms that the reattachment rate is independent of nucleotide conditions.

### Solution microtubule on-rates are also faster for kinesin-2 than kinesin-1

Because microtubule binding by a motor is inherently a bimolecular process, the first-order k_reattach_ parameter can be thought of as the product of a second-order microtubule on-rate multiplied by the effective local concentration of tubulin binding sites. Importantly, the scaffold, attachment, and coiled-coil domains are identical for the kinesin-1 and kinesin-2 assemblies used; hence the effective local [tubulin] should be identical for the two motors. In contrast, due to sequence differences in the microtubule binding domains and kinetic differences in their hydrolysis cycles, k_on_^Mt^ is expected to differ between kinesin-1 and kinesin-2. To test whether the different reattachment rates result from differences in the motor domains, we carried out stopped-flow experiments using the ADP analogue 2’(3’)-O-(N-methylanthraniloyl)adenosine 5’-diphosphate (mantADP) to measure the bi-molecular binding rate (k_on_^Mt^) for kinesin-1 and kinesin-2 (**Fig. 2D**). When motors incubated in mantADP are flushed against microtubules, microtubule binding triggers release of mantADP by the motor, which generates a decrease in mant fluorescence (**Fig. S2 E, F**). The process involves a sequential process of microtubule binding followed by mantADP release; hence, at saturating [Mt] the observed rate represents the mantADP off-rate of the microtubule-bound motor, whereas at limiting [Mt] the observed rate represents the on-rate for microtubule binding, k_on_^Mt^ (25). At each [Mt], fluorescence traces were fit by first-order exponentials (**Fig. S2 E, F**). The observed rate constants were then plotted as a function of [Mt] and fit with a line to obtain k_on_^Mt^ of 1.1±0.05 μM-1s^-1^ for kinesin-1 and 4.6 ±0.10 μM-1s^-1^ for kinesin-2 (**Fig. 2D, E**). Thus, the 3.6-fold higher k_reattach_ measured for kinesin-2 in the scaffold experiment matches the 4.2-fold higher k_on_^Mt^ for kinesin-2 in solution.

### High-resolution tracking reveals fast detachment/reattachment kinetics of kinesin-2

In order to measure detachment and reattachment events directly, we used high resolution single-molecule tracking to measure the time-dependent position of one kinesin in a two-motor pair attached to a DNA scaffold (**Fig. 3 A, B**). A kinesin-1 or -2 with a single motor domain biotinylated and tagged with a 30-nm gold nanoparticle (18, 22) was attached to one end in the scaffold, and an unlabeled motor was attached to the other (**Fig. 3B**). Example traces of Kin1-Kin1 and Kin2-Kin2 pairs are shown in **Fig. 3A**. Given that only one motor domain of one kinesin is labelled and the motors walk in a hand-over-hand manner, we expected to see low-variance ∼16 nm steps when the labelled kinesin was engaged with microtubule, higher-variance ∼8 nm steps when the labelled kinesin was not engaged with the microtubule, and large, abrupt positional changes when switching between these two configurations (**Fig. 3B**). We indeed observed these phenomena (**Fig. 3A**) among other features of note: (1) kinesin-1 spent longer durations with higher variance than kinein-2, as expected for their different reattachment rates, (2) newly reattaching kinesins landed both in front of and behind the currently engaged kinesin, and (3) kinesins commonly reattached to different protofilaments of the microtubule (as seen by positional changes perpendicular to the direction of motion).

**Figure 3:**
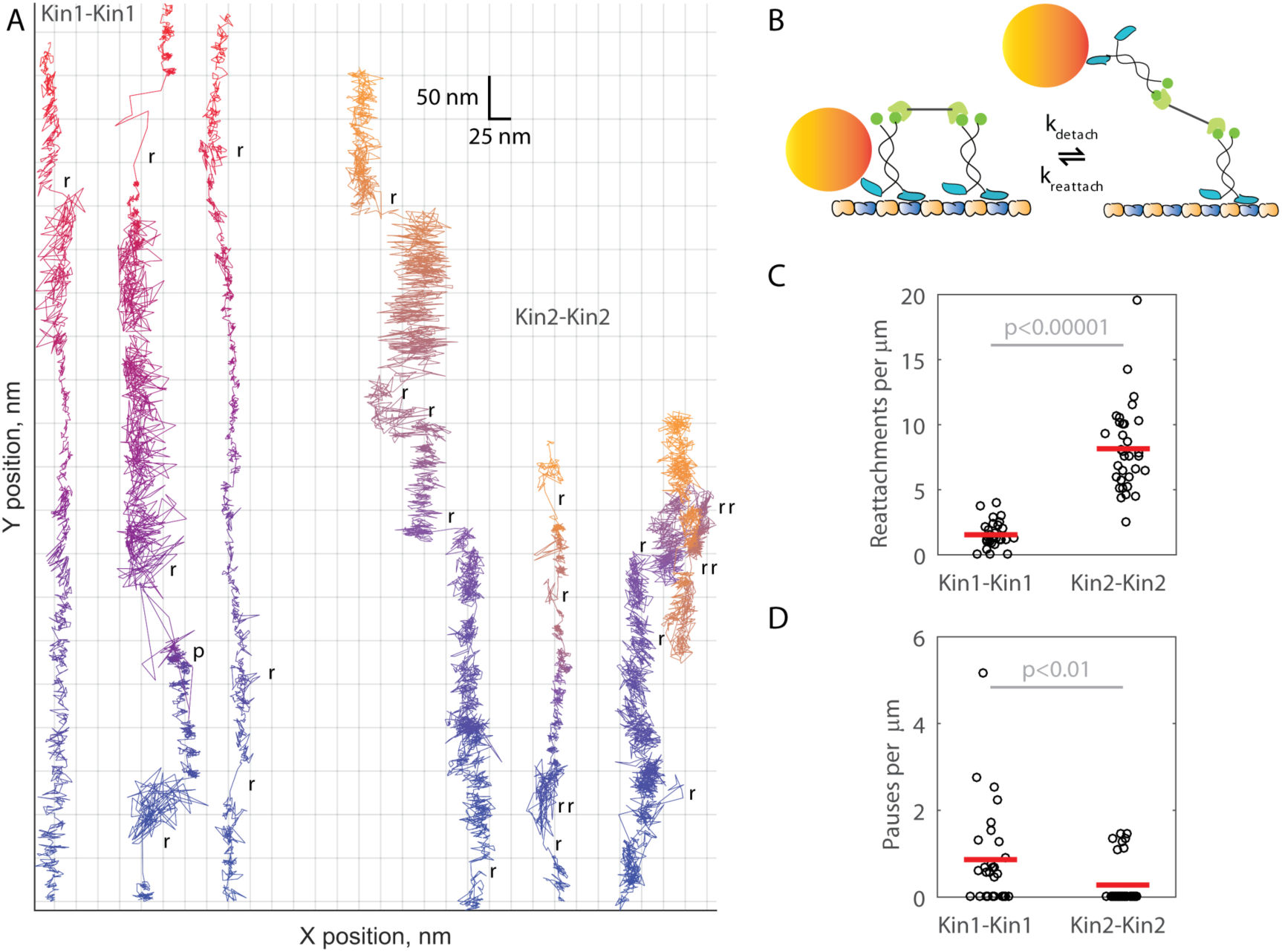
High resolution single-molecule tracking reveals that kinesin-2 reattaches more often and pauses less often than kinesin-1. (A) Example 1,000 frames per second traces of Kin1-Kin1 (blue-red) and Kin2-Kin2 (blue-yellow) pairs with a single motor domain of one motor tagged with a 30-nm gold nanoparticle (shown in diagram in (B)). Time information is encoded in color (see **Fig. S4** for the same data displayed as position versus time). Of note are abrupt positional changes that intersperse normal stepping, indicating reattachment events, and areas of high versus low variance, indicating whether one or two motors, respectively, are engaged with the microtubule. Scored rebinding events (**r**) and pauses (**p**) are highlighted on each trace. (C) Kin2-Kin2 pairs reattach more often than Kin1-Kin1 pairs. Reattachments were scored as jumps >40 nm in the Y position (parallel to the microtubule) or >15 nm in the X position (sidesteps). Kin1-Kin1 pairs reattached 1.54±0.19 times, while Kin2-Kin2 pairs reattached 8.16±0.58 times per micron traveled (mean±SEM; N=29 and N=33 traces, respectively, with plot showing one point per trace and mean values as red bars). A 2-sample T-test indicated that the difference in reattachment frequency was significant (P<0.00001). (D) Kin1-Kin1 pairs pause more often than Kin2-Kin2 pairs. Pauses were scored as instances of no positional change lasting longer than 10 step durations (137 ms for Kin1 and 410 ms for Kin2). Kin1-Kin1 pairs paused 0.86±0.21 times per micron traveled (mean±SEM, N=29 traces), while Kin2-Kin2 paused 0.28±0.09 times per micron traveled (mean±SEM, N=33 traces). All data shown, mean values shown as red bars. A Mann-Whitney U-test indicated that the difference in pausing frequency was significant (P<0.01).

To quantify the data, we scored detachment-reattachment events as positional jumps >40 nm (five tubulin lengths) parallel to the microtubule or >15 nm perpendicular to the microtubule. We observed that Kin2-Kin2 pairs reattached 5-fold more frequently per micron travelled than Kin1-Kin1 pairs (8.16 vs 1.54 reattachments/micron, respectively; **Fig. 3C**), in agreement with the reattachment rates in ADP (**Fig. 2)**. We also scored the pausing frequency, defined as the frequency the scaffold became stuck in a single position for more than 10 step-time durations. Kin1-Kin1 pairs paused 3-fold more frequently per micron travelled than Kin2-Kin2 pairs (0.86 vs 0.28 pauses/micron, respectively; **Fig. 3D**). These measurements are consistent with previous work that found kinesin-1 detachment is less sensitive to force than kinesin-2 (6, 7) and, together with the reattachment data, paint a picture of kinesin-1 being a fast, stable, but stubborn partner and kinesin-2 being a slow, vacillating, but adaptable partner in multi-motor transport.

### Kinesin-2 motors undergo fast detach/reattach cycles during multi-motor transport

To understand coordination between kin1 and kin2 motors during multi-motor transport, we measured the run length of kin1-kin2 pairs. Interestingly, despite the fact that kin2 has a shorter single-motor run length and a two-fold slower unloaded velocity than kin1, the run length of kin1-kin2 pairs, 2.18 ± 0.39 μm (**Fig. 4A**), was longer than for two kin1 motors 1.62 ± 0.23 μm (**Fig. 4B**). Thus, the faster reattachment rate of kin2 appears to be the key feature that enhances the multi-motor run length in motor pairs.

**Figure 4:**
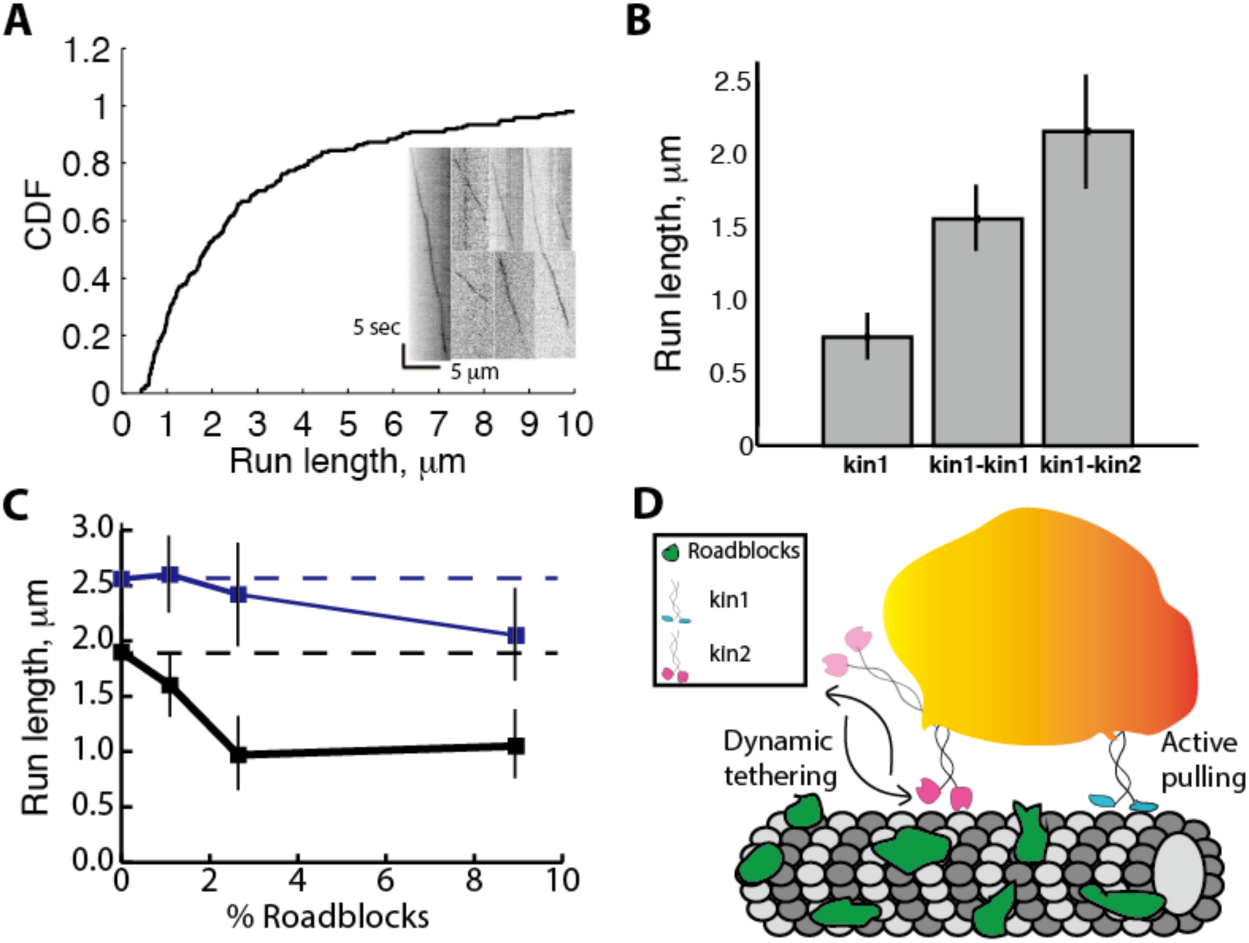
Cargo-bound kinesin-2 motors undergo fast detachment/reattachment to facilitate longer run lengths and avoid roadblocks. (A) Run length distributions and kymographs (inset) for Kin1-Kin2 pairs in 3 mM ATP. Mean kin1-kin2 run length was 2.18±0.39 μm (mean±95% confidence interval, N=199). See **Fig. S5A** for details of assembly. (B) Run lengths of single kinesin-1, kin1-kin1 pairs and kin1-kin2 pairs. Single kinesin-1 and kin1-kin1 run length are from **Fig. 1B**. (C) Run lengths of kin1-kin1 (black) and kin2-kin2 (blue) pairs on crowded microtubules. Dashed lines are run lengths without roadblocks for comparison. Roadblock concentrations are defined as the fraction of biotinylated tubulin in the microtubules with bound neutravidin. Run lengths are presented as mean±95% confidence intervals for between 25 and 117 measurements at each condition. See **Fig. S5B-D** for raw data. (D) Dynamic tethering model of kinesin-2 motors during intracellular cargo transport. In multi-motor assemblies, kinesin-2 motors (pink) will rapidly detach and reattach to the microtubule, while kinesin-1 motors (blue) will tend to remain bound to the microtubule and act as the primary force generators. This dynamic tethering of cargo to microtubules by kinesin-2 facilitates long distance transport and helps cargos navigate crowded microtubules.

To test the ability of motor multi-motor assemblies to avoid roadblocks such as MAPs we bound neutravidin to microtubules containing varying fractions of biotinylated tubulin and compared run lengths. Consistent with their fast detachment/reattachment kinetics, kin2-kin2 pairs were less affected by roadblocks than kin1-kin1 pairs (**Fig. 4C, S5C, D**). Thus kinesin-2 motors, despite moving slower and having a shorter unloaded run length and greater sensitivity of detachment to load, are able to coordinate their activities to achieve long multi-motor run lengths and navigate crowded microtubules.

## DISCUSSION

In cells, kinesin-1 and kinesin-2 each transport specific cargo, but they also colocalize on a subset of vesicles, suggesting that they also carry out coordinated transport (2, 3). In the present work, we show that kinesin-2 motors, despite being less processive than kinesin-1, enhance multi-motor run lengths to a greater degree and enable navigation of crowded microtubules. This behavior emphasizes the importance of motor reattachment rates on multi-motor transport.

### Fast reattachment is an inherent motor property

Despite the observed functional differences between kinesin-1 and kinesin-2, the specific amino acid sequences in kinesin-2 that confer faster microtubule rebinding kinetics property are not clear. For kinesin-3, the high initial microtubule binding rate is a result of its loop 12 domain, which has six positively-charged residues compared to only one for kinesin-1 (30). However, the kinesin-2 (KIF3A) loop 12 is nearly identical to kinesin-1, with the exception of having one less negatively charged residue (25). Similarly, the ADP off-rate upon microtubule binding is fast for both kinesin-1 and kinesin-2 (18, 25), suggesting that the probability of tight binding following collision with a microtubule is similar for the two motors. One possibility is that the fast microtubule on-rate of kinesin-2 is related to the motor’s propensity to remain associated with the microtubule in its weakly-bound state (25).

An important finding from comparing the measured bimolecular on-rates to the first order reattachment rates is that the effective local tubulin concentration is ∼30-fold lower than predicted from simple geometry considerations. This can be seen by considering that the reattachment rate is equal to the bimolecular on-rate multiplied by the effective local tubulin concentration, k_reattach_ = k_on_^Mt^ * [tubulin]. The predicted local tubulin concentration based on the motor-scaffold geometry can be calculated as follows. If the tethered motor searches a hemispherical volume with a radius of ∼ 100 nm that contains six protofilaments (the top half of the microtubule), the tubulin concentration in this hemisphere is 125 μM (see **Fig. S3E**). Multiplying this concentration by the measured k_on_^Mt^ = 4.6 μM^-1^S^-1^ for kinesin-2 (**Fig. 3E**) results in a predicted k_reattach_ of >500 s^-1^, compared to the 16 s^-1^ measured value. The source of this discrepancy is not clear.

One intriguing finding from comparing the present work to previous studies of defined pairs of kinesin-1 motors linked through DNA (31, 32) or protein scaffolds (33, 34) is that the run length enhancement from adding a second motor is consistently quite small, ranging from 1.3-fold to 2.5-fold (31-33). Furthermore, previous work showed that when the length and rigidity of a DNA linker connecting the motors were systematically varied over a large range, there was very little effect on run length (31), consistent with the motor reattachment rate being relatively insensitive to the specific properties of the linker that connects the two motors. The reattachment rate of kinesin-1 motors attached to membranes was previously estimated at 4.7 s^-1^, matching our estimate, despite the very different geometries (15). The enhancement of run length by kinesin-2 observed here suggests that the microtubule binding properties of the motor domains play the dominant role in motor reattachment kinetics rather than the specific geometry of the scaffold. Understanding the tethered diffusion that leads to these observed motor reattachment rates is an important area for future investigations.

### Kinesin-1 and -2 motors are tuned for different cellular roles in multi-motor transport

The fast kinesin-2 reattachment rate measured here provides resolution for the earlier work that showed detachment of heterotrimeric kinesin-2 depends strongly on load (6-9). The present work establishes that the propensity of kinesin-2 to detach under load is balanced by rapid reattachment, which results in the motor actually spending most of its time bound to the microtubule in a multi-motor system. The present work also provides an explanation for the earlier finding that purified neuronal vesicles have both kinesin-1 and kinesin-2 motors bound, despite the fact that they move at two-fold different speeds (3). We propose a model in which kinesin-1 is an “active puller” that generates the force needed for transport while kinesin-2 serves as a “dynamic tether” (**Fig. 4D**). This dynamic tethering serves first to maintain association of the cargo with the microtubule when kinesin-1 motors detach, and second to enable cargos to navigate along microtubules crowded with MAPs and other impediments without becoming stalled. This tethering activity may explain a body of previous work on bidirectional transport that found that inhibiting either kinesin or dynein alone diminishes transport in both directions (2). If this tethering activity of kinesin-2 also helps to maintain association of the cargo with the microtubule while dynein is pulling, then inhibiting the motor may diminish this tethering activity and thus diminish dynein-driven transport. Because kinesin-3 is able to diffuse on microtubules and has fast initial microtubule attachment kinetics (1, 33), this behavior is predicted to extend to kinesin-3 as well.

In conclusion, the present work presents a method for quantifying the motor reattachment rate in multi-motor assemblies and demonstrates that k_reattach_ is four-fold faster for kinesin-2 than kinesin-1. The prediction of fast binding/unbinding kinetics for kinesin-2 is directly demonstrated using high-resolution tracking of one motor, a technique that can be extended to more complex multi-motor geometries. Finally, we show that kinesin-2 motor pairs more effectively navigate crowded microtubules. This work provides important foundational pillars for quantitatively understanding the complex motor dynamics underlying bidirectional transport of vesicles in cells.

## ACKNOWLEDGEMENTS

We are thankful to David Arginteanu for help with protein purification, the Grischuk lab for providing GBP plasmids, Rizal Hariadi and the Sivaramakrishnan lab for assistance with DNA origami, and Hancock lab members for helpful discussions. This work was funded by NIH R01 GM076476 and GM121679 to W. O. H.

